# Interrogating the Topological Robustness of Gene Regulatory Circuits

**DOI:** 10.1101/084962

**Authors:** Bin Huang, Mingyang Lu, Dongya Jia, Eshel Ben-Jacob, Herbert Levine, Jose N. Onuchic

## Abstract

One of the most important roles of cells is performing their cellular tasks properly for survival. Cells usually achieve robust functionality, for example cell-fate decision-making and signal transduction, through multiple layers of regulation involving many genes. Despite the combinatorial complexity of gene regulation, its quantitative behavior has been typically studied on the basis of experimentally-verified core gene regulatory circuitry, composed of a small set of important elements. It is still unclear how such a core circuit operates in the presence of many other regulatory molecules and in a crowded and noisy cellular environment. Here we report a new computational method, named random circuit perturbation (RACIPE), for interrogating the robust dynamical behavior of a gene regulatory circuit even without accurate measurements of circuit kinetic parameters. RACIPE generates an ensemble of random kinetic models corresponding to a fixed circuit topology, and utilizes statistical tools to identify generic properties of the circuit. By applying RACIPE to simple toggle-switch-like motifs, we observed that the stable states of all models converge to experimentally observed gene state clusters even when the parameters are strongly perturbed. RACIPE was further applied to a proposed 22-gene network of the Epithelial-to-Mesenchymal transition (EMT), from which we identified four experimentally observed gene states, including the states that are associated with two different types of hybrid Epithelial/Mesenchymal phenotypes. Our results suggest that dynamics of a gene circuit is mainly determined by its topology, not by detailed circuit parameters. Our work provides a theoretical foundation for circuit-based systems biology modeling. We anticipate RACIPE to be a powerful tool to predict and decode circuit design principles in an unbiased manner, and to quantitatively evaluate the robustness and heterogeneity of gene expression.

## Introduction

State-of-the-art molecular profiling techniques[1–4] have enabled the construction or inference of large gene regulatory networks underlying certain cellular functions, such as cell differentiation[5,6] and circadian rhythm[7,8]. However, it remains a challenge to understand the operating principles of these regulatory networks and how they can robustly perform their tasks, a prerequisite for cell survival. Mathematical and computational systems biology approaches are often applied to quantitatively model the dynamic behaviors of a network[9–20]. Yet, quantitative simulations of network dynamics are usually limited due to several reasons. First, a proposed network might contain inaccurate or missing regulatory genes or links, and modeling an incomplete network might produce inaccurate predictions. Second, kinetic parameters for each gene and regulatory interaction, which are usually required for quantitative analyses, are difficult to be obtained directly for all of them from *in vivo* experiments[21]. To deal with this problem, network parameters are either inferred from existing data [22,23] or educated guesses, an approach which could be time-consuming and error-prone. This approach is hard to extend to very large gene networks due to their complexity.

Alternatively, a bottom-up strategy has been widely used to study the regulatory mechanisms of cellular functions. First, one performs a comprehensive analysis and integration of experimental evidence for the essential regulatory interactions in order to construct a core regulatory circuit, typically composed of only a small set of essential genes. The core gene circuit is then modeled either by deterministic or stochastic approaches with a particular set of parameters inferred from the literature. Due to the reduced size of the systems and the inclusion of data derived directly from the literature, the bottom-up approach suffers less from the above-mentioned issues. Examples of the bottom-up approach include the modeling of biological process such as Epithelial-to-Mesenchymal transition (EMT)[24–26], cell cycles[27,28], and circuit design in synthetic biology, such as genetic toggle switch[29], and repressilator[30].

Due to the success of these and other circuit-based modeling studies, we hypothesize that a core circuit module should emerge from a complex network and dictate the decision-making process. It is reasonable to assume that a large gene network could be decomposed into a core gene circuit and a peripheral part with the residual genes. The core would then be the driving force for the network dynamics and should be robust against cell-to-cell variability and extrinsic fluctuations in stimuli arising from cell signaling, while the peripheral genes would act to regulate the signaling status for the core circuit and probably also enhance the robustness of the core dynamics by introducing redundancy. This scale-separation picture is consistent with ideas such as the existence of master regulators and network modularity[31,32].

On the basis of this conceptual framework, we developed a new computational method, named *<u>ra</u>*ndom *<u>ci</u>*rcuit *<u>pe</u>*rturbation (RACIPE), for modeling possible dynamic behaviors that are defined by the topology of a core gene regulatory circuit. In RACIPE, we focus the modeling analysis on the core circuit and regard the effects of the peripheral genes and external signaling as random perturbations to the kinetic parameters. In contrast to traditional modeling methods[33], RACIPE generates an ensemble of mathematical models, each of which has a different set of kinetic parameters representing variations of signaling states, epigenetic states, and genetic backgrounds (including cells with genetic mutations leading to disease). Here we randomize the model parameters by one or two orders of magnitude and utilize a specially designed parameter sampling scheme (details in Methods) to capture the key role of circuit topology. This random field approach allows the inclusion of the contributions from the peripheral genes to the network dynamics and the evaluation of their roles in modulating the functions of the core circuit. From the *in silico* generated data, we apply statistical analysis to identify the most probable features within all of the models, a process which can uncover the most robust functions of the core circuit. It is worth-noting that RACIPE is unique in the way it utilizes perturbation and the integration of statistical tools, compared to the traditional parameter sensitivity analysis[34–38] and the previous studies on random circuit topology[39,40].

In the following, we will first describe in detail the RACIPE method, and then present the results of applying RACIPE to several simple standalone circuit motifs and also coupled toggle switch motifs. In addition, we will show the application of RACIPE to a 22-component network for the decision-making core of the Epithelial-to-Mesenchymal Transition (EMT). We will see that RACIPE is capable of identifying accessible gene states via statistical analysis of the *in silico* generated data, from which we can further decode the design principles and evaluate the robustness of the core gene circuit. We therefore expect RACIPE to be a powerful tool to analyze the dynamic behavior of a gene network and to evaluate the robustness and accuracy of proposed network models.

## Methods

We developed a new computational method, namely *ra*ndom *<u>ci</u>*rcuit *<u>pe</u>*rturbation (RACIPE), for modeling a gene network. The procedure of RACIPE is illustrated in Fig. 1. The input of RACIPE is the topology of the core circuit under study, which can be constructed according to either the literature, interaction databases (e.g. Ingenuity pathway analysis (IPA®, QIAGEN Redwood City, www.qiagen.com/ingenuity), KEGG[41], GO[42]), or computational methods[43]. From the circuit topology, we establish a set of mathematical equations for the time evolution of the levels of all the genes. We then generate an ensemble of models where the parameters from the rate equations are sampled by a carefully designed randomization procedure (see below for details) so that these kinetic models can capture the behavior of the circuits under different conditions. Each model is subject to standard analysis to discover possible dynamics of the circuit (Fig. 1B). The dynamics could converge to a stable steady state, a stable oscillation, or chaotic behavior. To find all possible behaviors for a gene network, we typically choose many different sets of initial conditions (randomly sampled on a logarithmic scale) and numerically solve the rate equations for each case. This ODE-based method is particularly useful for identifying all the distinct stable steady states for a multi-stable system. The procedure is repeated for many times to collect sufficient data for statistical analysis. In particular, for a multi-stable system, the RACIPE method generates a large amount of simulated gene expression data, which can be further analyzed by biostatistics tools (Fig. 1C). RACIPE is also compatible with other types of modeling methods such as the stochastic approach, but this is not discussed in this study. In the following, we will illustrate RACIPE in the context of a multi-stable gene circuit by deterministic analysis.

**Fig 1.**
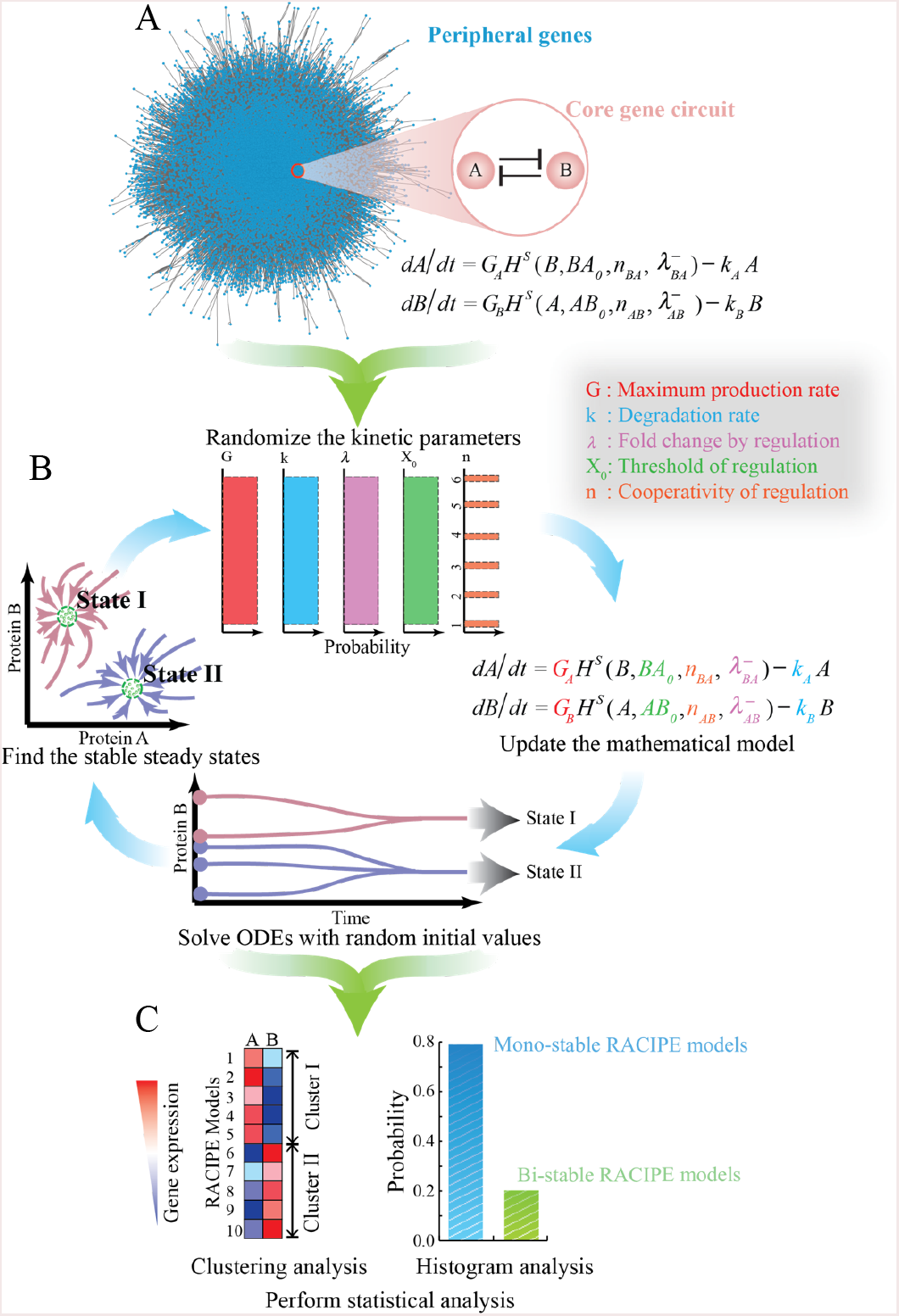
Schematics of the random circuit perturbation (RACIPE) method. (A) The gene regulatory network for a specific cellular function is decomposed into two parts - a core gene circuit modeled by chemical rate equations and the other peripheral genes whose contribution to the network is regarded as random perturbations to the kinetic parameters of the core circuit; **(B)** RACIPE generates an ensemble of models, each of which is simulated by the same rate equations but with randomly sampled kinetic parameters. For each model, multiple runs of simulations are performed, starting from different initial conditions, to identify all possible stable steady states; **(C)** The *in silico* gene expression data derived from all of the models are subject to statistical analysis.

As an example, we start with the deterministic rate equations for a toggle switch circuit (Fig. 2) with mutually inhibitory genes A and B. The kinetic model takes the form:

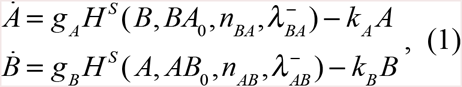

where *A* and *B* represent the expression levels of gene A and B respectively. *g*_*A*_ and *g*_*B*_ are the basal production rates (the production rate for the gene without any regulator bound to the promoter). *k*_*A*_ and *k*_*B*_ are the innate degradation rates. Regulation of gene B expression by A is formulated as a non-linear shifted Hill function 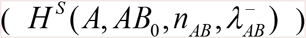, defined as 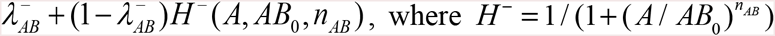 is the inhibitory Hill function, *AB*_0_ is the threshold level for A, *n*_*AB*_ is the Hill coefficient of the regulation, 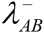 is the maximum fold change of the B level caused by the inhibitor A 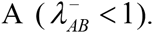 In the case of an activator, the fold change is represented by 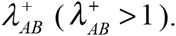 The inhibitory regulation of gene A by gene B can be modeled in an analogous way.

**Fig 2.**
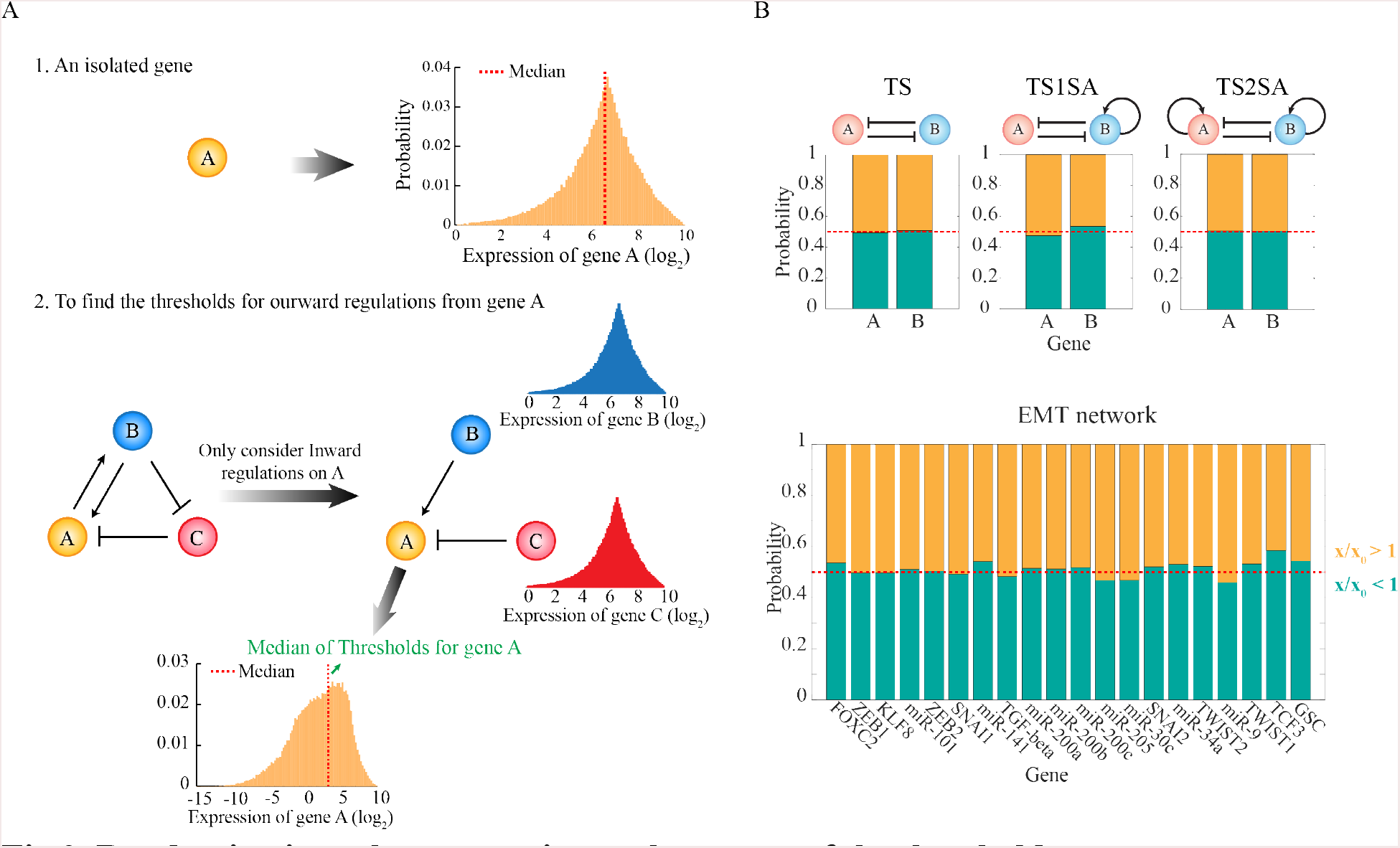
Randomization scheme to estimate the ranges of the threshold parameters. (A) Schematics of the procedure to estimate the ranges of the threshold parameters, so that the level of a regulator has 50% chance to be above or below the threshold level of each regulatory link (“half-functional rule”). First, for a gene A without any regulator, the RACIPE models are generated by randomizing the maximum production rate and the degradation rate according to S1 Table. The distribution of A level is obtained from the stable steady state solutions of all the RACIPE models (top left panel, yellow histogram). Second, for a gene A in a gene circuit, the distribution of A level is estimated only on the basis of the inward regulatory links (i.e. the B to A activation and the C to A inhibition in the bottom left panel). The distributions of the levels of the inward regulators B and C are assumed to follow the same distributions as a gene without any regulator (bottom left panel, blue and red distribution); the threshold levels for these inward links are chosen randomly from (0.02M to 1.98M), where M is the median of their gene expression distributions. Finally, the distribution of A level is obtained by randomizing all the relevant parameters. That includes the levels of B and C, the strength of the inward regulatory links, the maximum production rate and the degradation rate of the A, and the threshold for any regulatory link starting from A is chosen randomly from (0.02M to 1.98M), where M is the median level of the new distribution of A level (orange in the bottom panel). The same procedure is followed for all other genes. **(B)** Tests on several simple toggle-switch-like circuit motifs and the Epithelial-to-Mesenchymal Transition (EMT) circuit show that the “half-functional rule” is approximately satisfied with this randomization scheme. For each RACIPE model, we computed the ratio (x/x_0_) of the level of each gene X at each stable steady state (x) and the threshold (x_0_) for the outward regulations from gene X. The yellow region shows the probability of x/x_0_ > 1 for all the RACIPE models, and the green region shows the probability of x/x_0_ < 1.

In RACIPE, randomization is performed on all five types of circuit parameters: two of them are associated with each gene, including the basal production rate (*g*) and the degradation rate (*k*); and three of them are associated with each regulatory link, including the maximum fold change of the gene expression level (*λ*), the threshold level of the regulation (*X*_0_) and the Hill coefficient (n).

Our parametric randomization procedure is designed to ensure that the models can represent all biologically relevant possibilities. In detail, the Hill coefficient *n* is an integer selected from 1 to 6, and the degradation rate *k* ranges from 0.1 to 1 (See S1 Table for the explanation of the units). Here each parameter is assigned by randomly picking values from either a uniform distribution or some other distributions, for example the Gaussian distribution. The fold change *λ*^+^ ranges from 1 to 100 if the regulatory link is excitatory, while *λ*^-^ was varied from 0.01 to 1 if the regulatory link is inhibitory. Note that for the latter case, a probability distribution (e.g. a uniform distribution) is sampled for the inverse of *λ*^-^, i.e. 1 / *λ*^-^, instead of *λ*^-^ itself. By doing so, we make sure that the mean fold change is about 0.02, instead of ~ 0.5. The choice of such a wide range of *λ* values ensures the consideration of both strong and weak interactions.

In addition, two assumptions are made in RACIPE to ensure that it generates a representative ensemble of models for a specific circuit topology. First, the maximum production rate of each gene should lie roughly within the same range (from 1 to 100 in this study, see S1 Table), as the maximum rate is determined by how fastest the transcriptional machinery can work. For a gene regulated by only one activator, the maximum production rate is achieved when the activator is abundant, and thus the basal production rate of the gene *g* = *G* / *λ*^+^. For a gene regulated by only one inhibitor, the maximum rate is achieved in the absence of the inhibitor, i.e. *g* = *G*. This criterion can be generalized to genes regulated by multiple regulators. Therefore, in practice, we directly randomize the maximum production rate (G) instead of the basal production rate (g), and then calculate the value of g according to the above criterion. The ranges of these parameters are summarized in details in S1 Table. The RACIPE randomization procedure allows a gene to have a relative expression ratio of up to 1,000 for two sets of parameters, even when it is not regulated by other genes.

Second, we also assume that, for all the members of the RACIPE model ensemble, each regulatory link in the circuit should have roughly equal chance of being functional or not functional, referred to as the *half-functional* rule. For example, in the case that gene A regulates gene B, all the parameters are selected in such a way that for the RACIPE ensemble, the level of A at the steady states has roughly 50% chance to be above and 50% chance to be below its threshold level. Otherwise, if the threshold level is too large or too small, the regulatory link is either not functional most time or constitutively active, thereby changing the effective circuit topology.

To achieve this, we estimate the range of the threshold levels by a mean-field approximation. For a regulatory link from gene A (regulator) to gene B (target), the threshold level *AB*_0_ can be estimated as follows. We first estimate the range of expression of gene A without considering any of its regulators. The A level without regulation satisfies

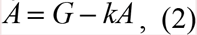

By randomizing both *G* and *k* by the aforementioned protocol (S1 Table), we generate an ensemble of random models, from which we obtain the distribution of the steady state levels of gene *A* (Fig. 2A). To meet the half-functional rule, the median of the threshold level should be chosen to be the median of this distribution. When gene A is regulated by some other genes (i.e. its upstream regulators), we estimate the median threshold level by considering the inward regulators of A, and assume that the levels of all these regulators (*e.g.* gene B, C *etc.*) follow the same distribution as an isolated gene (top right panels in Fig. 2A section 2). We set the threshold of every inward regulation to be 0.02M to 1.98M, where M is the median of the distribution of an isolated gene. With all of the information, we can again generate a new ensemble of models, from which we calculate the distribution of gene A (bottom panel in Fig. 2A section 2) and its median. For every target gene regulated by the gene A, we select the threshold levels of the regulations in the range from 0.02M to 1.98M, where M is the above obtained median level of gene A. The same approach is used to estimate the threshold levels of the other genes. It is worth-noting that this self-consistent strategy works quite well for the cases we have tested (Fig. 2B) according to the *half-functional* rule.

In the following, we will first demonstrate the application of RACIPE to some simple toggle-switch-like motifs, then to a set of motifs of coupled toggle-switch circuits, and eventually to a more complex EMT transcription regulatory network. For each case, we will illustrate how we can utilize an ensemble of RACIPE models to identify the dynamic behavior of a gene circuit.

## Results

### RACIPE as an unbiased method to predict robust gene states for a gene circuit

We first tested RACIPE on several basic toggle-switch-like circuit motifs (Fig. 3A). These circuit motifs are considered to be some of the main building blocks of gene regulatory networks[44]. A genetic toggle switch (TS), composed of two mutually inhibitory genes, is commonly considered to function as a bi-stable switch - it allows two stable gene states, each of which is characterized by dominant expression of one gene. The TS has been shown to be a central piece of decision-making modules for cell differentiation in several incidences[45–47].

**Fig 3.**
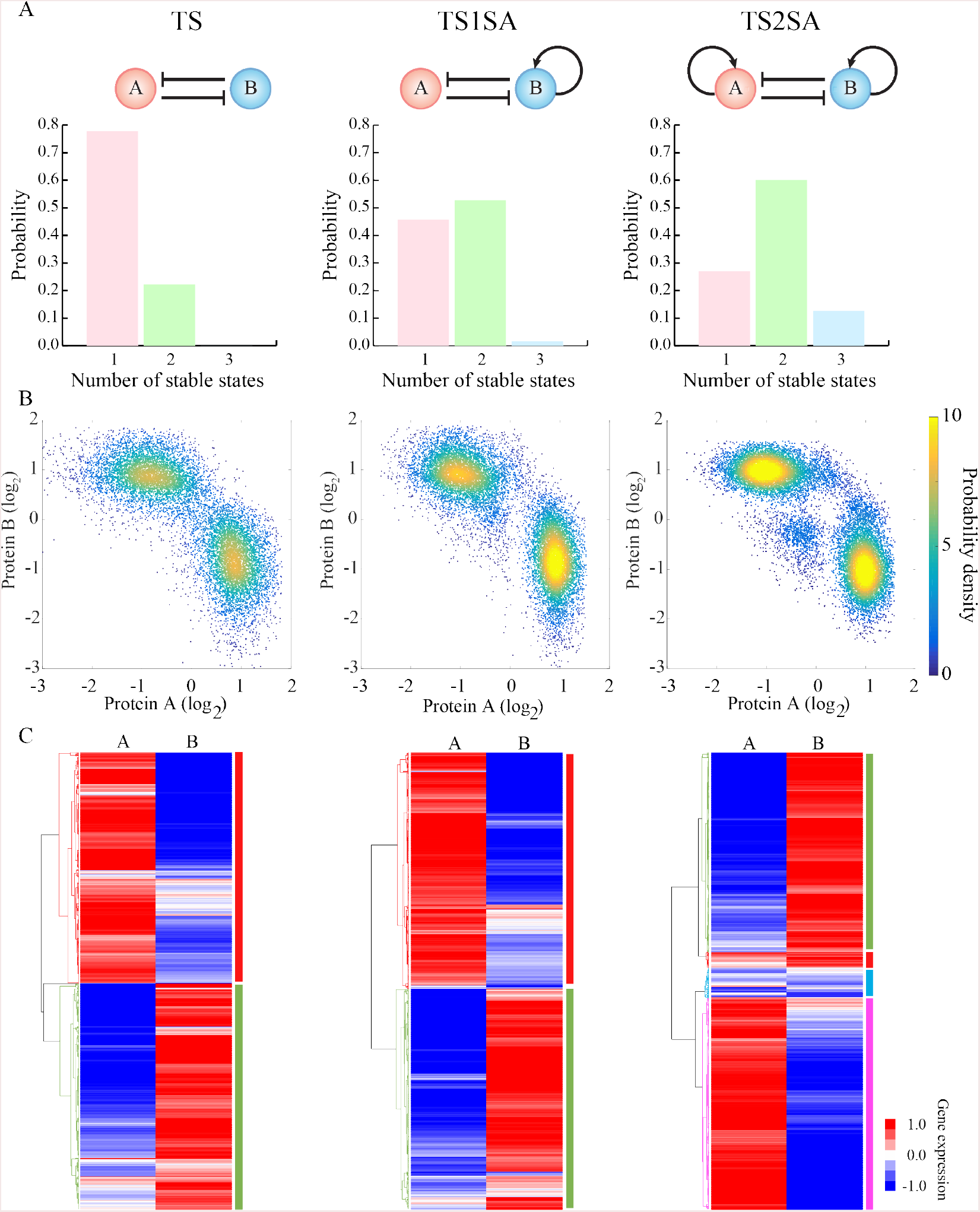
RACIPE identifies robust features of toggle-switch-like motifs. RACIPE was tested on three circuits – a simple toggle-switch (TS, top left) which consists of genes A and B that mutually inhibit each other (solid lines and bars), a toggle-switch with one-sided self-activation (TS1SA) which has an additional self-activation link on gene B, and a toggle-switch with two-sided self-activation (TS2SA) which has additional self-activation links on both genes. **(A)** Probability distributions of the number of stable steady states for each circuit. (B) Probability density maps of the gene expression data from all the RACIPE models. Each point represents a stable steady state from a model. For any RACIPE model with multiple stable steady states, all of them are shown in the plot. **(C)** Average linkage hierarchical clustering analysis of the gene expression data from all the RACIPE models using the Euclidean distance. Each column corresponds to a gene, while each row corresponds to a stable steady state from a model. The analysis shows that the gene expression data could be clustered into distinct groups, each of which is associated with a gene state, as highlighted by different colors on the right of the heatmaps.

Here we apply RACIPE to the TS motif. We created an ensemble of 10,000 models (Fig. 3A) and we observed that about 20% of models allow two coexisting stable steady states (bi-stability), while the remainder allow only one steady state (mono-stability). The observation that only a small fraction of TS models work as a bi-stable system is consistent with a previous study[39]. Next, we tested RACIPE on a toggle switch with an extra excitatory auto-regulatory link acting on only one of the genes (a toggle switch with one-sided self-activation, or TS1SA). The circuit motif now has ~ 50% chances of being bi-stable, much larger than the original TS motif. Interestingly, TS1SA also has ~1% chance of having three co-existing stable steady states (tri-stability), so it can potentially act as a three-way switch[44]. Hence, the RACIPE analysis suggests that TS1SA is more robust than TS for functioning as a switch. Moreover, adding excitatory auto-regulatory links on both sides of the TS motif (TS2SA) further increases the likelihood of bi-stability to ~60%, and meanwhile dramatically increases the likelihood of tri-stability to ~13%. This suggests that TS2SA has more of an ability than these other motifs to function as a three-way switch. Indeed, TS2SA has been proposed to be the core decision-making motif for several cell differentiation processes, and many of these processes exhibit multi-stability[45,46]. Thus, statistical analysis of the ensemble of random models generated by RACIPE can identify the most robust features of a circuit motif.

Another way to utilize RACIPE is to evaluate the possible gene expression patterns of a circuit motif. We can construct a large set of *in silico* gene expression data, consisting of the gene expression levels of the circuit at every stable steady state for each RACIPE model. In the dataset, the column corresponds to the genes and the rows corresponds to stable steady states. For a RACIPE model with multiple stable steady states, we enter the data in multiple rows. The expression dataset takes a form similar to typical experimental microarray data, and so it can be analyzed using common bioinformatics tools. For each of the above two-gene cases, we visualized the expression data by a scatter plot of the levels of the two genes (Fig. 3B). Surprisingly, despite large variations in the circuit parameters across the RAICPE model ensemble, the expression data points converge quite well into several robust clusters. For example, the TS circuit data has two distinct clusters, where one has a high expression of gene A but a low expression of gene B and the other vice versa. The TS2SA circuit has not only the above two clusters but also an additional cluster with intermediate expression of both genes. These patterns have also been observed in previous experimental[29] and theoretical[44,45,48] studies of the same circuits. Interestingly, if we only include models with a fixed number of stable states (e.g. restrict the ensemble to mono-stable models, or bi-stable models), a similar pattern of clusters can still be observed (Fig. S1). These clusters represent distinct patterns of gene expression that the circuit can support, so we will refer to these clusters as “gene states”. These gene states are robust against large perturbations of circuit parameters because the circuit topology restricts possible gene expression patterns. RACIPE in a sense takes advantage of this feature to interrogate the circuit so that we can unbiasedly identify the robust gene states. Since these states may be associated with different cell phenotypes during cell differentiation or cellular decision-making processes, RACIPE can be especially helpful in understanding the regulatory roles of the circuit during transitions among different states.

These simple cases demonstrate the effectiveness of RACIPE in revealing generic properties of circuit motifs. Recall that our basic hypothesis is that the dynamic behaviors of a circuit should be mainly determined by circuit topology, rather than a specific set of parameters. The rich amount of gene expression data generated by RACIPE allows the application of statistical learning methods for the discovery of these robust features. For example, as shown in Fig. 3C, we applied unsupervised hierarchical clustering analysis (HCA) to the RACIPE gene expression data, and again we identified similar gene state clusters for each circuit. Notably, the predictions of these gene states by RACIPE should be robust against different sampling distributions and different ranges of kinetic parameters. To verify this, we tested on the TS circuit versions of RACIPE created with three different distributions (uniform, Gaussian and exponential distribution) and three different ranges of parameters (Fig. 4). Even though the precise shape of gene states appears to be slightly different for the different cases, the number and the locations of these gene states are consistent (Fig. 4). For the cases with exponential distribution, when the range of the parameters is reduced, the mean decreases as well; therefore, the two gene states become closer (Fig. 4). We also found that the expansion of the parameter ranges still results in similar gene states (Fig. S2).

**Fig 4.**
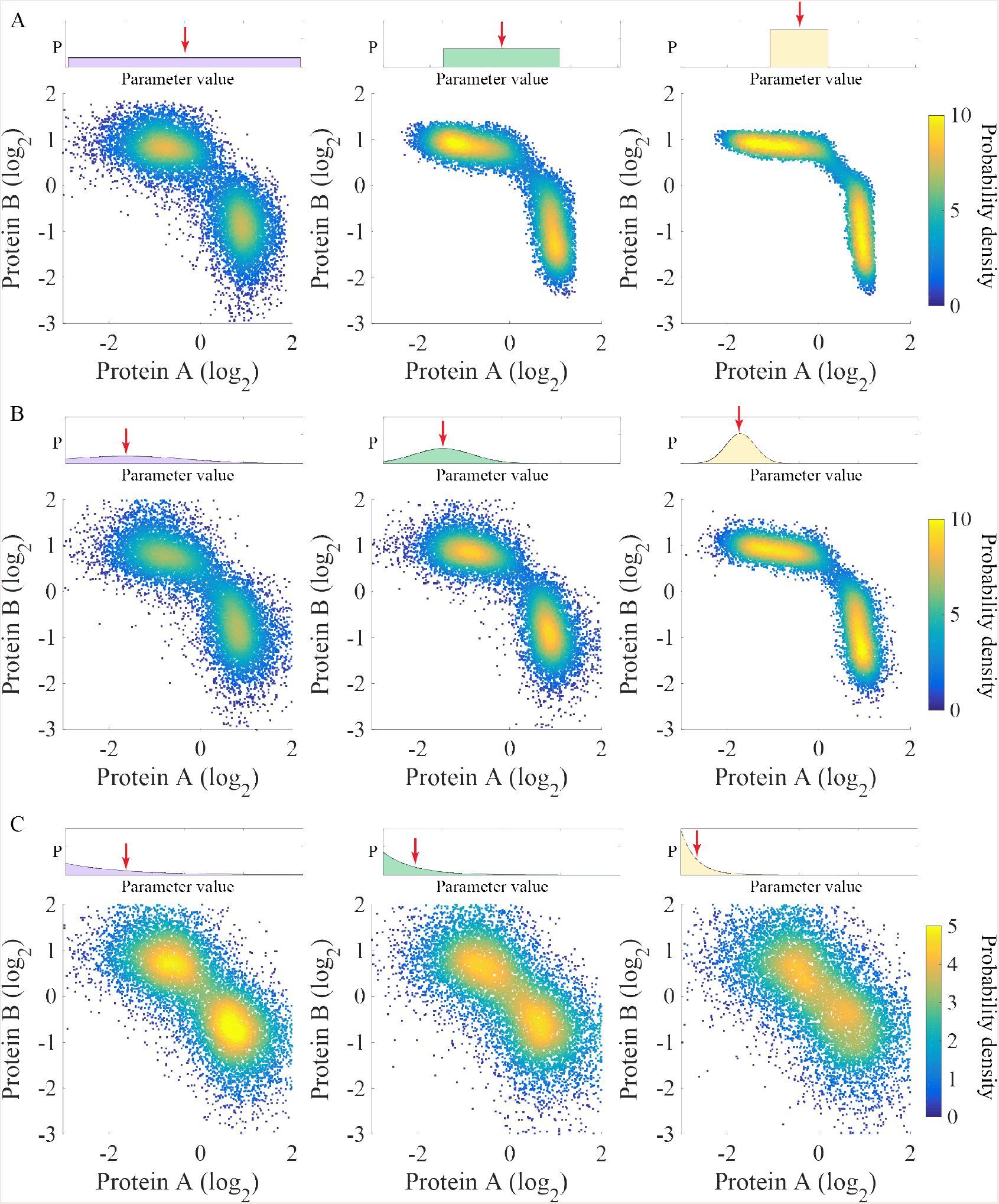
The gene states of the toggle-switch motif are robust against different types of distributions used to sample the parameters. (A) Uniform distributions in three different ranges were used to sample the kinetic parameters of the RACIPE models. The top panels show the range of the distribution (left panel: the full range; middle panel: half; right panel: 1/4). The bottom panels show the probability density maps of the gene expression data from all the RACIPE models. Similarly, panels (B) and (C) show the use of a Gaussian distribution and an exponential distribution, respectively. For the Gaussian distribution (B), its standard deviation was shrunk by a factor of two from left to right. For the Exponential distribution (C), its mean was reduced by a factor of two from left to right.

### The application of RACIPE to coupled toggle-switch motifs

To evaluate the effectiveness of RACIPE on larger circuits, we further applied the method to circuits with two to five coupled toggle-switch (CTS) motifs (Fig. 5). Different from the above simple circuit motifs, the gene expression data obtained by RACIPE for these CTS motifs are now high-dimensional; thus in the scatter plot analysis we projected these data onto the first two principal components by principal component analysis (PCA). For each circuit, we observed distinct gene states from PCA for the RACIPE models (Fig. 5A). More interestingly, the number of gene states found via PCA increases by one each time one more toggle switch is added to the circuit. Moreover, we applied HCA to the gene expression data, from which we identified the same gene states as from PCA (Fig. 5B). At this stage, we can also assign high (red circles), intermediate (blue circles) or low expression (black circles) to each gene for every gene state. Unlike in Boolean network models, the assignment in RACIPE is based on the distribution of expression pattern from all the models in the ensemble (Fig S3-S4).

**Fig 5.**
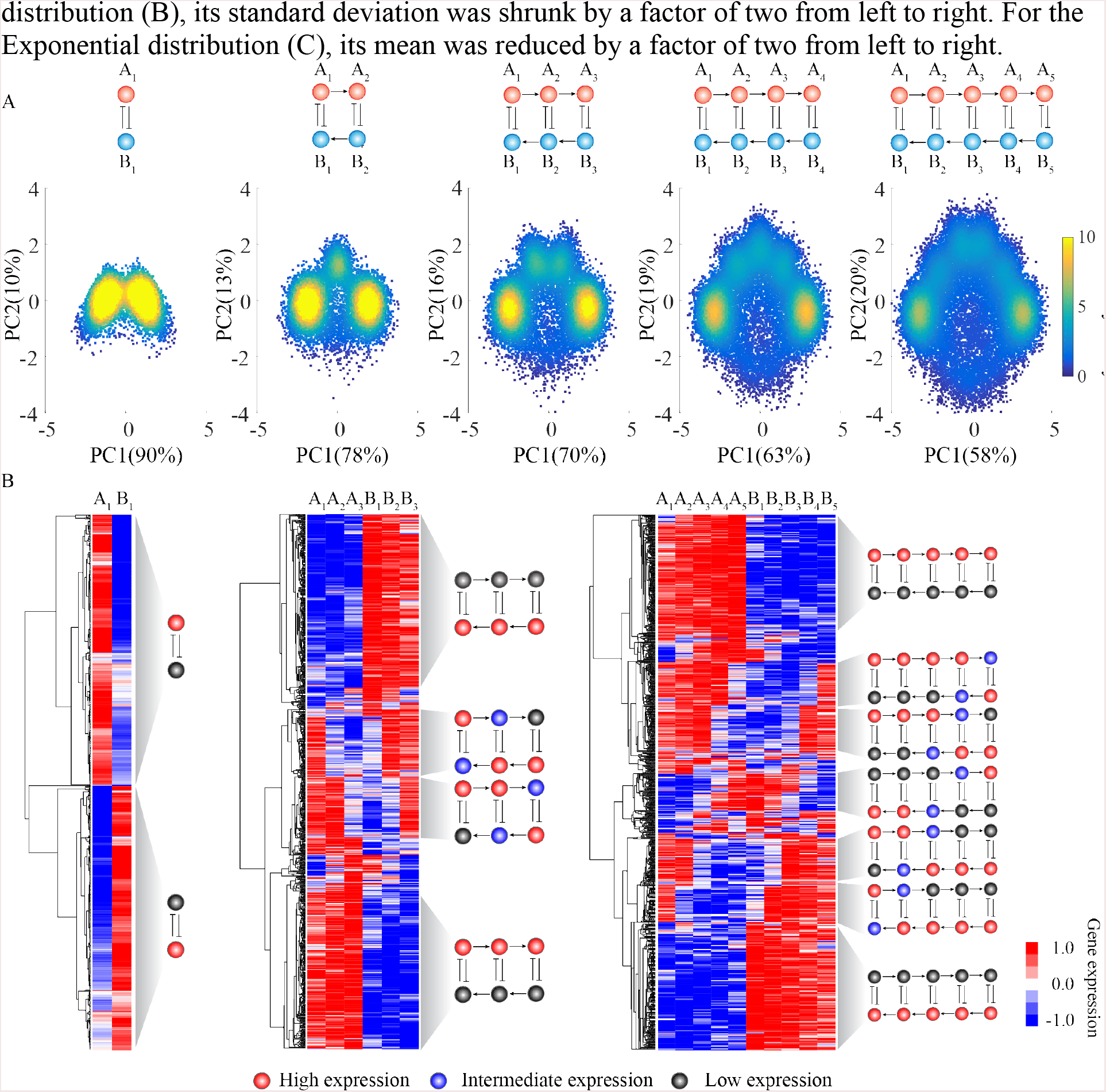
Application of RACIPE to coupled toggle-switch circuits. RACIPE was tested on coupled toggle-switch circuits, as illustrated at the top of the figure. **(A)** 2D probability density map of the RACIPE-predicted gene expression data projected to the 1^st^ and 2^nd^ principal component axes. **(B)** Average linkage hierarchical clustering analysis of the gene expression data from all the RACIPE models using the Euclidean distance. Each column corresponds to a gene, while each row corresponds to a stable steady state. The clustering analysis allows the identification of several robust gene states, whose characteristics were illustrated as circuit cartoons to the right of the heatmap. The expression levels of each gene in these gene states are illustrated as low (grey), intermediate (blue), or high (red, see SI for the definition).

We can easily understand the meaning of each gene state. In each case, the rightmost cluster in the scatter plot (Fig. 5A) corresponds to the topmost cluster in the heatmap (Fig. 5B), a state where all the A genes have high expression and all the B genes have low expression. Similarly, the leftmost cluster in the scatter plot corresponds to the bottommost cluster in the heatmap. These two clusters are the most probable ones, and represent the two extreme states of the coupled toggle switch network. As also illustrated in the scatter plots, for circuits with additional toggle switches, these two states separate from each other and the circuit now allows intermediate states. By closely examining these intermediate states, we found that they (from top to bottom) correspond to a cascade of flips of the state of each consecutive toggle switch. This explains why we observe one additional gene state every time we include an additional toggle-switch motif. In addition, intermediate expression levels were frequently observed for genes lying in the middle toggle-switch motifs, instead of those at the edge. The tests on CTS circuits demonstrate again the power of RACIPE in identifying robust features of a complex circuit.

### The application of RACIPE to the EMT circuit

The above examples were used for illustrative purposes and do not immediately reflect any actual biological process. In our last example, we apply RACIPE to a more realistic case, the decision-making circuit of EMT (Fig. 6). EMT is crucial for embryonic development, wound healing and metastasis[49], a major cause for 90% cancer-related deaths[50]. Cells can undergo either a complete EMT to acquire mesenchymal phenotype or partial EMT to attain hybrid E/M phenotype[51,52], which maintains both E and M traits. Transitions among the Epithelial (E), Mesenchymal (M) and hybrid epithelial/mesenchymal (E/M) phenotypes have been widely studied either experimentally or theoretically[52]. Here, we utilized data from the literature and Ingenuity Pathway Analysis (see details in SI) to construct a core gene regulatory circuit model of EMT (Fig. 6A), which contains 13 transcriptional factors (TFs), 9 microRNAs (miRs) and 82 regulatory links among them. Among the gene components, we have two biomarkers - CDH1 and VIM - that are commonly used to distinguish different phenotypes during EMT, and one signaling gene TGF-β. The circuit is a much-extended version of several previous EMT models[24,25], which consist of only four gene families and one input signal. It is similar in terms of scale to a recently proposed Boolean model of EMT[53], but as stressed here our models allow for continuous expression levels.

**Fig 6.**
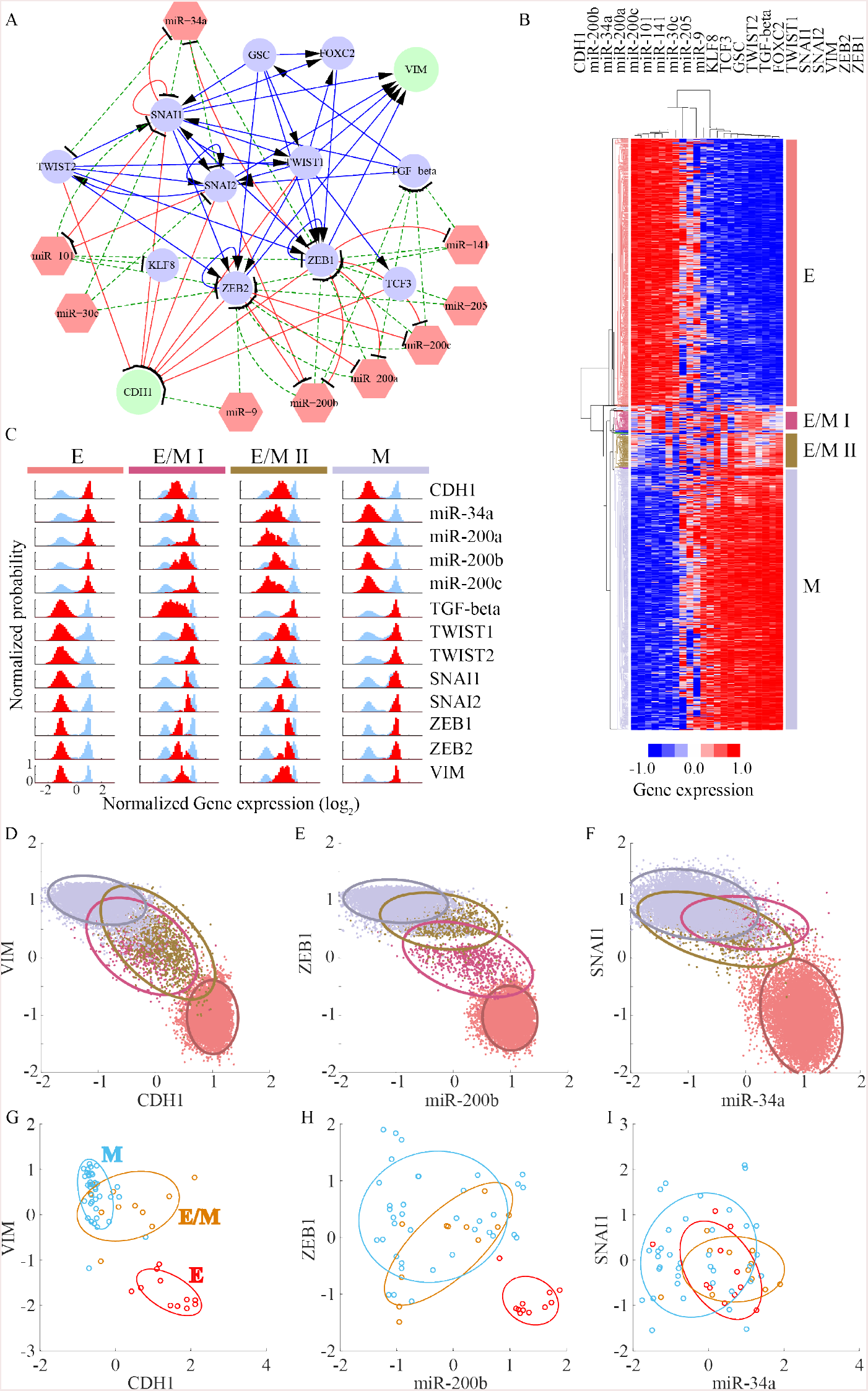
RAICPE identifies multiple EMT cell states from gene network analysis. **(A)** A proposed Epithelial-to-Mesenchymal Transition (EMT) circuit is constructed according to the literature; the circuit consists of 13 transcriptional factors (circles), 9 microRNAs (red hexagons) and 82 regulatory links. The blue solid lines and arrows represent transcriptional activations the red solid lines and bars represent transcriptional inhibition, and the green dashed lines and bars stand for translational inhibition. Two readout genes CDH1 and VIM are shown as green circles while the other transcriptional factors are shown in blue. **(B)** Average linkage hierarchical clustering analysis of the gene expression data from all the RACIPE models using the Euclidean distance. Each column corresponds to a gene, and each row corresponds to a stable steady state. Four major gene states were identified and highlighted by different colors. According to the expression levels of CDH1 and VIM, the four gene states were associated with epithelial (E in red), mesenchymal (M in grey) and two hybrid epithelial/mesenchymal (E/M I in purple and E/M II in brown) phenotypes. **(C)** The gene expression distribution of each gene state. The gene expression distribution of each gene for all of the RACIPE models is shown in blue, while that for each gene state is shown in red (50 bins are used to calculate the histogram of each distribution). For clarity, each distribution is normalized by its maximum probability. Each row represents a gene and each column represents a gene state. **(D-F)** Gene expression data were projected to either CDH1/VIM, miR-200b/ZEB1, or miR-34a/SNAI1 axes. Different gene states are highlighted by the corresponding colors and enclosed by the ellipses. **(G-I)** Transcriptomics data from the NCI-60 cell lines were projected to either CDH1/VIM, miR-200b/ZEB1, or miR-34a/SNAI1 axes. The NCI-60 cell lines have been grouped into E, E/M and M phenotypes according to the ratio of the protein levels of CDH1 and VIM. Different gene states are highlighted by the corresponding colors and enclosed by the ellipses.

For simplicity, we modeled the EMT circuit with the same approach as above, i.e. all the genetic components were coupled with Hill functions, typical of transcriptional control. This may not be completely accurate for translational regulation by microRNA (miR), but we leave this complication for future study. Even with this simplification, RACIPE can already provide insightful information of the EMT regulation. Consistent with what we learned from the test cases, unsupervised HCA of the RACIPE gene expression data can reveals distinct gene states (Fig. 6B). Here there are four such states. We can map these gene states to different cell phenotypes possible during EMT - an E phenotype with high expression of the miRs, low expression of TFs, and CDH1^HI^VIM^LO^; a M phenotype with low expression of the miRs, high expressions of TFs, and CDH1^LO^VIM^HI^; and two hybrid E/M phenotypes with intermediate expression of both miRs, TFs and CDH1/VIM. The E/M I state lies closer to the E state, and the E/M II state lies closer to the M state. More intriguingly, we found SNAI1 and SNAI2 become highly expressed in the E/M I phenotype while ZEB1 and ZEB2 are not fully expressed until the E/M II or the M phenotype (Fig. 6C), which is a possibility supported by recent experimental results[25].

Moreover, RACIPE can help to find genes of similar function and filter out less important genes in the core circuit. As shown in Fig. 6B, genes are grouped into two major clusters according to their expression levels throughout all the RACIPE models - miRs/CDH1 and TFs/VIM. The former genes are highly expressed mainly in E phenotypes while the latter are highly expressed in M phenotypes. We also found three microRNAs (miR-30c, miR-205 and miR-9) to be randomly expressed in the RACIPE models, indicating these three microRNAs are less important to these EMT phenotypes. From the topology of the circuit, we see that these three microRNAs lack feedback regulation and act solely as inputs.

A typical approach taken in cell biology is to use two biomarkers to identify cells of different states in a mixed population by fluorescence-activated cell sorting (FACS). To mimic the analysis, we projected the gene expression data of the RACIPE models onto the two axes of important genes, as shown in the scatter plots in Fig. 6D-F. In all of the three cases, the E and the M phenotypes can be distinguished. However, for the hybrid phenotypes, the E/M I and the E/M II states overlap in the CDH1-VIM plot (Fig. 6D). These two hybrid phenotypes can be separated more easily in the ZEB1-miR200b plot (Fig. 6E). In the SNAIL1-miR34a plot (Fig. 6F), however, the two E/M states overlap with the M state. The theoretical prediction that the SNAIl1-miR34a axis is less efficient at distinguishing the states is supported by transcriptomics data from the NCI-60 cell lines[54] (Fig. 6G-I). We see here that either VIM-CDH1 or the ZEB1-miR200b axes are indeed better than the SNAIL1-miR34a axes in separating different EMT phenotypes. Our results are also consistent with our previous theoretical finding that ZEB1 is more crucial than SNAIL1 in the decision-making of EMT[25].

## Discussion

Recently, the rapid development of genomic profiling tools has allowed the mapping of gene regulatory networks. Yet, it remains a challenge to understand the operating mechanisms and the design principles of these networks. Conventional computational modeling methods provide insightful information; however, their prediction power is usually limited by the incompleteness of the network structure and the absence of reliable kinetics. To deal with these issues, we have developed a new computational modeling method, called RACIPE, which allows unbiased predictions of the dynamic behaviors of a complex gene regulatory circuit. Compared to traditional methods, RACIPE uniquely generates an ensemble of models with distinct kinetic parameters. These models can faithfully represent the circuit topology and meanwhile capture the heterogeneity in the kinetics of the genetic regulation. By modeling the dynamics of every RACIPE model, we can utilize statistical analysis tools to identify the robust features of network dynamics. We have successfully tested RACIPE on several theoretical circuit motifs and a proposed core Epithelial-to-Mesenchymal Transition (EMT) gene regulatory circuit. In each circuit, RACIPE is capable of predicting the relevant gene states and providing insights into the regulatory mechanism of the decision-making among gene states.

Unlike other methods that utilize randomization strategies to explore the parameter sensitivity for gene circuit[34–37], RACIPE adopts a carefully designed sampling strategy to randomize circuit parameters over a wide range. Instead of looking for the sensitivity of the circuit function to parameter variations, we focused on uncovering conserved features remaining in the ensemble of RACIPE models. This was carried out by statistical learning methods such as hierarchical clustering analysis. We showed the power of RACIPE to predict the robust gene states for a circuit with a given topology. However, the rich data generated by RACIPE can be further analyzed for more hidden information. Moreover, it is easy to implement gene modifications such as knockdown or overexpression treatments with the RACIPE method to learn the significance of each gene or link in the circuit. Therefore, RACIPE provides a new way to model a gene circuit without knowing the detailed circuit parameters.

Another parameter-independent approach people often use for gene circuit modeling is Boolean network model[55], which digitalizes the gene expression into on and off states and uses logic functions to describe the combinatorial effects of regulators to their targets. Compared with the Boolean network model, RACIPE is a continuous method, so it is not restricted to the on and off values. Instead, RACIPE enables us to find the intermediate levels of gene expressions beyond the on and off states, as we showed in Fig. 5B and Fig. 6C. From the ensemble of RACIPE models, we can predict the expression distribution of the gene, which can be directly compared with experimental expression data. In addition, in RACIPE, we not only obtain *in silico* gene expression data, but we also know the kinetic parameters for each model. From these parameter data, we can directly compare the parameter distributions for different gene states, from which we can learn the driving parameters that are responsible for the transitions among the states.

To conclude, here we have introduced a new theoretical modeling method, RACIPE, to unbiasedly study the behavior of a core gene regulatory circuit under the presence of intrinsic or extrinsic fluctuations. These fluctuations could represent different signaling environments, epigenetic states, and/or genetic backgrounds of the core circuit and can cause cell-cell heterogeneity in a population. By approximating these fluctuations as variations of the model parameters, RACIPE provides a straightforward way to understand the heterogeneity and to explain further how gene circuits can perform robust functions under such conditions. Moreover, RACIPE uniquely generates a large set *in silico* expression data, which can be directly compared with experimental data using common bioinformatics tools. RACIPE enables the connection of traditional circuit-based bottom-up approach with profiling-based top-down approach. We expect RACIPE to be a powerful method to identify the role of network topology in determining network operating principles.

## Acknowledgements

This work was supported by National Science Foundation (NSF) Center for Theoretical Biological Physics (NSF PHY-1427654) and NSF grant MCB-1214457. H.L. and J.N.O. are also supported as CPRIT (Cancer Prevention and Research Institute of Texas) Scholar in Cancer Research of the State of Texas at Rice University. M.L. was also supported by a training fellowship from Keck Center for Interdisciplinary Bioscience Training of the Gulf Coast Consortia (CPRIT Grant RP140113). This work was supported in part by the Big-Data Private-Cloud Research Cyberinfrastructure MRI-award funded by NSF under grant CNS-1338099 and by Rice University.

## Author contributions

B.H. and M.L. developed the method, and performed the simulations, analyzed the data, and drafted the manuscript. D.J analyzed the EMT experimental results. E.B.J., H.L. and J.N.O. supervised the work, participated in the analysis of the data, and revised the manuscript. All authors (other than E.B.J.) read and approved the final manuscript.

